# Neuronal inhibition of the autophagy nucleation complex extends lifespan in post-reproductive *C. elegans*

**DOI:** 10.1101/135178

**Authors:** Thomas Wilhelm, Jonathan Byrne, Rebeca Medina, Ena Kolundzic, Johannes Geisinger, Martina Hajduskova, Baris Tursun, Holger Richly

**Affiliations:** Laboratory of Molecular Epigenetics, Institute of Molecular Biology (IMB), Mainz, Germany, Ackermannweg 4, 55128 Mainz, Germany; Faculty of Biology, Johannes Gutenberg University, 55099 Mainz, Germany; Berlin Institute for Medical Systems Biology (BIMSB), Max Delbrück Center (MDC), Robert-Rössle-Straße 10, 13125 Berlin, Germany

**Author notes:** These authors contributed equally (Thomas Wilhelm, Jonathan Byrne).

**Keywords:** Antagonistic pleiotropy, *C. elegans*, BEC-1, autophagy, aging, neurodegeneration

## Abstract

Autophagy is a ubiquitous catabolic process, which causes cellular bulk degradation of cytoplasmic components and thereby regulates cellular homeostasis. Inactivation of autophagy has been linked with detrimental effects to cells and organisms. The antagonistic pleiotropy theory postulates that fitness promoting genes during youth are harmful during aging (Williams 1957). On this basis we examined genes mediating post-reproductive longevity using an RNA interference screen. From this screen we identified 30 novel regulators of post-reproductive longevity including *pha-4*. Through downstream analysis of *pha-4* we identify that genes governing the early stages of autophagy up until the stage of vesicle nucleation, such as *bec-1*, strongly extend both lifespan and healthspan. Further, our data demonstrates that the improvements in health and longevity are mediated through the neurons - resulting in reduced neurodegeneration and sarcopenia. We propose that autophagy switches from advantageous to harmful in the context of an age-associated dysfunction.

## Introduction

Aging represents the functional deterioration of an organism, which compromises fitness and predisposes for prevalent diseases such as cancer and neurodegeneration. Many genes that modulate aging have been identified in *C. elegans* through mutagenesis or RNAi screens. These screens inactivate genes across the whole lifespan from early development (Ni and Lee 2010; Klass 1983). Genes identified through these studies often work through a small set of common pathways, including the target of rapamycin (TOR) pathway and the insulin like signaling pathway (Ni and Lee 2010). However, genetic pathways specific to aging in older individuals remain largely undiscovered. The existence of alleles belonging to such pathways is predicted by the antagonistic pleiotropy (AP) theory of aging. This evolutionary-based theory states that strong natural selection early in life enriches alleles that mediate fitness, while simultaneously accumulating their harmful effects after reproduction, when natural selection is ineffectual (Williams 1957). Thus, some genes that reduce fitness when inhibited in young worms should conversely extend lifespan when inhibited in older worms. Two screens in late L4 worms identified AP genes critical to development (Curran and Ruvkun 2007; Chen et al. 2007). However, these screens were limited in their detection potential for AP genes, as they initiated RNAi early in life, at a point of maximal selection pressure (Fig. 1A).

**Fig. 1.**
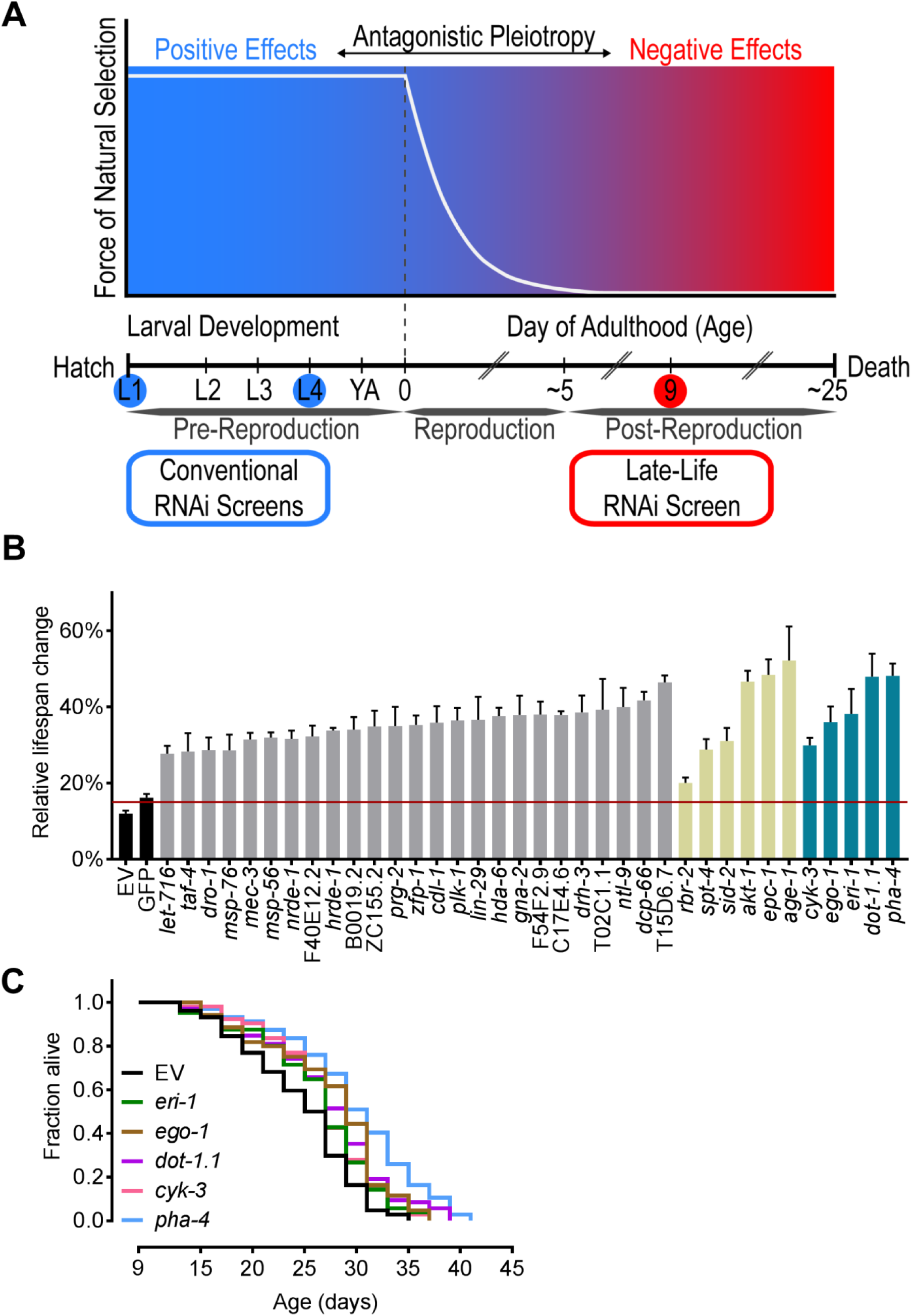
Screening for AP genes uncovers novel regulators of post-reproductive longevity. (A) Screening for antagonistic pleiotropic (AP) genes. The force of natural selection (white line) declines over age. AP genes have positive fitness effects (blue region) early in life and negative effects (red region) late in life. Inhibiting genes post-reproductively can identify novel regulators of longevity. Conventional RNAi screens’ initiation point (blue circles). Initiation point of our RNAi screen (red circle). (B) Potential longevity genes identified from the AP RNAi screen. The percentage of *rrf-3(pk1426)* worms alive at day 30 is shown. Controls: Empty vector (EV) and non-targeting GFP (black), novel longevity genes (grey), known longevity genes (khaki) and our top five candidate genes (blue). Red line indicates the threshold for consideration as a longevity candidate. Data represents the average of 4 replicates +/**-** the SEM (Supplemental Table 2). (C) Day 9 RNAi against our top candidate genes: *eri-1*, *dot-1.1*, *cyk-3*, *ego-1*, and *pha-4* extend mean treated lifespan (MTL). — All lifespans were *rrf-3(pk1426).* Days represent days post first egg lay. Lifespan statistics, Supplemental Table 4.

Autophagy is a conserved process that encloses cytoplasmic components and organelles in double membrane structures called autophagesomes, which then fuse with the lysosome where the contents are degraded and recycled (Mizushima et al. 1998). As such, autophagy is essential for proteostasis, whereby: long-lived, unfolded, misfolded, or damaged proteins are removed from circulation. Autophagy is required for normal development, health, and is crucial for the extended lifespan by reduced germline-, insulin-or TOR-signaling (Rubinsztein et al. 2011). Autophagy is predominantly cytoprotective, however, when autophagy becomes dysregulated, it is associated with adverse effects such as cell death or failure to clear aggregated proteins resulting in neurodegeneration (Komatsu et al. 2006). Autophagy has in fact been called a double edged sword in that it can be detrimental in certain disease contexts either being causal or exacerbating their pathologies (Shintani and Klionsky 2004). Yet, the role of autophagy in post-reproductive lifespan has not been investigated.

Here, we demonstrate a novel post-reproductive screening approach for the identification of unknown longevity genes functioning according to the AP model of aging (Fig. 1A). In our screen we identified 30 novel regulators of longevity, including the FOXA transcription factor *pha-4.* We find that PHA-4 inactivation increases lifespan through its role in autophagy via BEC-1. Importantly, the inhibition of *pha-4* or *bec-1* modulate lifespan conversely over aging fulfilling the criteria for AP. Our data indicates that global autophagy becomes dysfunctional with age and is blocked at the later steps of autophagosomal degradation. We demonstrate that post-reproductive inhibition of the VPS-34/BEC-1/EPG-8 autophagic nucleation complex, as well as its upstream regulators, strongly extend *C. elegans* lifespan. Further, we show that post-reproductive inhibition of *bec-1* mediates longevity specifically through the neurons. Our data suggests that suppression of early autophagy in aged worms results in improved neuronal integrity, contributing to enhanced global health and culminating in increased longevity.

## Results

### Screening for AP genes identifies regulators of post-reproductive longevity

To screen for potential AP genes that extend post-reproductive lifespan, we assembled an RNAi library targeting 800 gene-regulatory factors involved in chromatin or transcriptional regulation (Supplemental Table 1). These targets were chosen as disruption of such factors is often detrimental to organismal fitness, a requirement for AP. As we inhibit these genes late in life, we were concerned that decreased feeding rates in aged worms (Huang et al. 2004) would preclude robust RNAi phenotypes. Consequently, the screen was carried out in *rrf-3* mutant animals, which exhibit enhanced sensitivity to RNAi (Simmer et al. 2002). To generate the large numbers of post-reproductive and age-synchronized worms required for this screen, we developed a large-scale liquid culture system (Supplemental Fig. S1, A to D). This system allowed for screening in fertile worms, excluding the potential cofounding effects of germline removal or 5-fluoro-2’-deoxyuridine (FUdR) treatment. We conducted gene inactivation at day 9 by sorting worms into 24-well agar plates, which contained the RNAi feeding library. Longevity candidates were identified by the presence of live worms at day 32, when the majority of the control worms were dead. We identified 36 longevity genes (5% of the library), 30 of which have never previously been linked with lifespan extension (Fig. 1B and Supplemental Table 2). Examining the listed phenotypes for these new longevity genes in WormBase (version WS258), we found that 19 are good AP candidates owing to phenotypes associated with lethality, as well as with defects in development, reproduction, growth, and health (Supplemental Table 2). We validated the efficacy of our screen in identifying novel longevity regulators through conventional lifespan assays of our top five candidates (Fig. 1C). The observed extension of mean treated lifespan (MTL) for these candidates ranged from 12% (*eri-1*) to 33% (*pha-4*). We used the *rrf-3* strain for follow-up experiments unless stated otherwise.

### PHA-4 and BEC-1 exhibit AP and improve healthspan upon post-reproductive inhibition

We focused on PHA-4/FOXA as a prime AP candidate given its crucial role in development, targeting over 4000 genes (Zhong et al. 2010) and as its early inactivation is known to reduce lifespan (Sheaffer et al. 2008). Activation of *pha-4* induces autophagy and regulates longevity in diet-restricted and germline-less animals (Lapierre et al. 2011; Hansen et al. 2008). Additionally, the critical autophagy nucleation factor BEC-1 is a transcriptional target of PHA-4 (Zhong et al. 2010). In general, autophagy is predominantly associated with its positive effects on health and longevity (Rubinsztein et al. 2011). Given the strong association of *pha-4* and autophagy, we determined if autophagy could have a pleiotropic role in post-reproductive longevity.

As previously shown in young worms (Lapierre et al. 2011), we demonstrated that both key autophagy genes *bec-1* and *unc-51* remain regulated by PHA-4 in aged worms (Supplemental Fig. S2A). Further, GFP::LGG-1 foci formation, which marks autophagosomal membranes, was strongly reduced upon *pha-4* and *bec-1* inactivation at day 9 (Supplemental Fig. S2, B to E). By inactivating *bec-1* from day 9, we tested if PHA-4 mediates its longevity effect through BEC-1. This intervention caused a 63% MTL extension, twice that of *pha-4* inactivation (Fig. 2A and Supplemental Fig. S2F). As BEC-1 acts downstream of PHA-4 in the autophagic cascade (Lapierre et al. 2011), the differences in MTL extension were unanticipated. The reduced lifespan effect of PHA-4 inactivation could be due to its broad spectrum of target genes (Zhong et al. 2010), some of which might have detrimental effects when dysregulated. In support, simultaneous inactivation of *pha-4* and *bec-1* at day 9 caused significant reductions in *bec-1* lifespan extension (Fig. 2B).

**Fig. 2.**
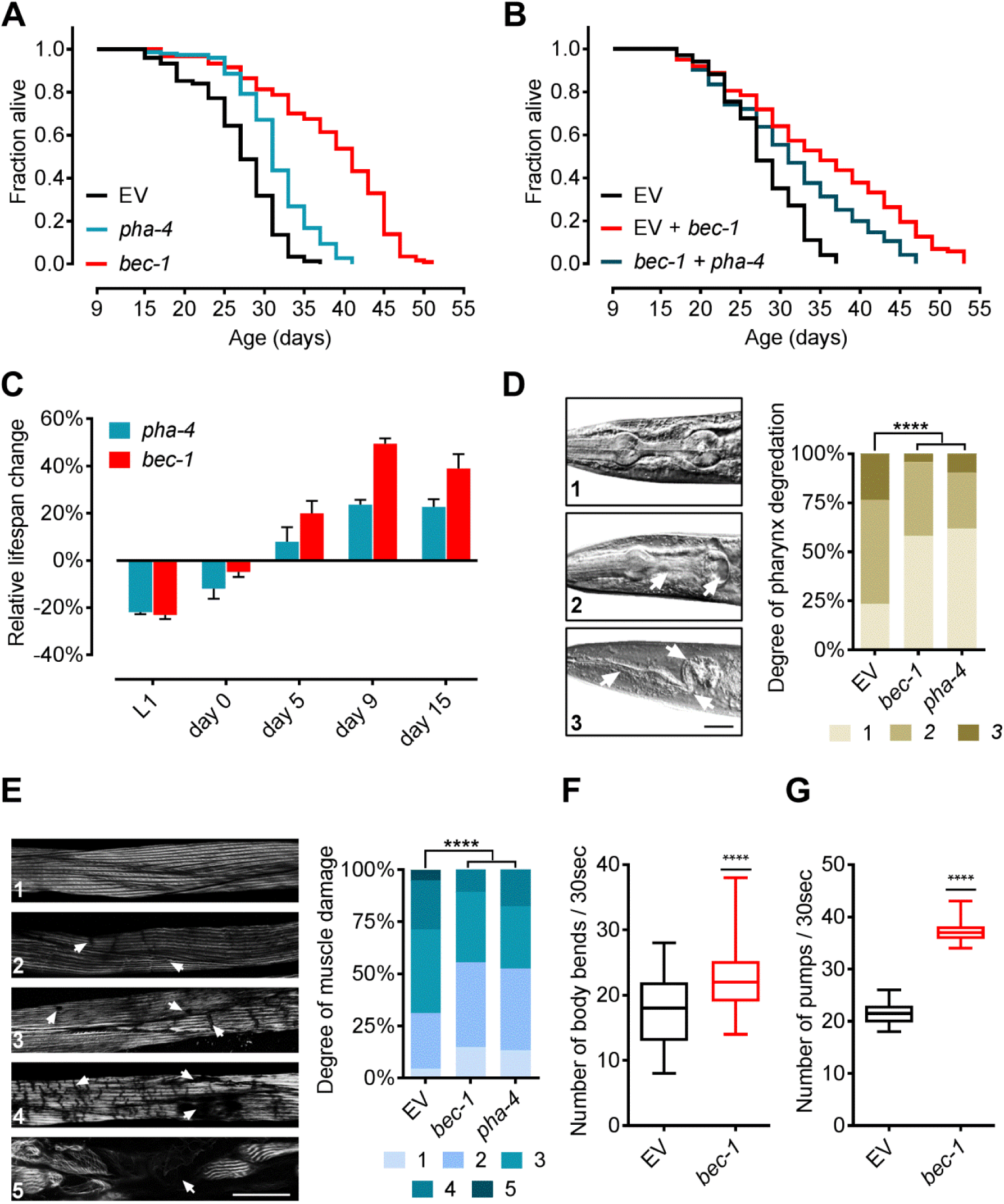
*pha-4* and *bec-1* extend lifespan in an AP manner and improve post-reproductive healthspan. (A) Day 9 RNAi against *bec-1* extends MTL by twice that of *pha-4*. (B) Inactivation of *pha-4* at day 9 in combination with *bec-1* reduces MTL extension compared to *bec-1* knockdown alone. (C) Inhibition of *pha-4* and *bec-1* at different times across life shows differing effects depending on the RNAi initiation time point. Depicted is the relative percentage change in MTL +/**-** SEM compared to control. Knockdown validation (Supplemental Fig. S3, D and E). (D) Day 9 RNAi treated worms were scored at day 20 for pharynx degradation on a three point scale (N=90). Damage of the corpus, isthmus and terminal bulb (white arrows). Scale bar, 30 μm. (**** = p<0.0001). (E) Day 9 RNAi treated worms were scored at day 20 for phalloidin stained muscle damage on a five point scale (N=120). Sites of muscle damage (white arrows). Scale bar, 30 μm (**** = p<0.0001). (F) Day 20 quantification of body movement, measured by body bend counts, in *rrf-3(pk1426)* worms. Data is depicted as a box plot with min to max values. RNAi against *bec-1* was initiated at day 9 (N=50) (**** = p<0.0001). (F) Day 20 quantification of pharynx pumping rate in *rrf-3(pk1426)* worms. Data is depicted as a box plot with min to max values. RNAi against *bec-1* was initiated at day 9 (N=50) (**** = p<0.0001). — All lifespans were *rrf-3(pk1426).* Days represent days post first egg lay. Lifespan statistics, tables S4 and S5.

The AP hypothesis predicts different longevity effects based on the gene inactivation time-point. Consequently, we examined if *pha-4* and *bec-1* knockdowns across lifespan exhibit matching AP effects. We observed a decrease in MTL in larvae and young adult worms, transitioning into a neutral, then positive effect during aging for both genes (Fig. 2C). We determined if the observed lifespan extension of both genes translates into an improved healthspan. Inactivation of *pha-4* or *bec-1* at day 9 caused the preservation of a youthful pharynx and muscle structure (Fig. 2, D and E). In addition, *bec-1* knockdown at day 9 significantly improved mobility and pharynx pumping (Fig. 2, F and G). These findings suggest that late-life inactivation of both genes extend lifespan and improve healthspan. Collectively, these results imply that *pha-4* inactivation mediates lifespan extension through autophagy via BEC-1. As an integral autophagic component, we proceeded with BEC-1 to further investigate the role of autophagy in the observed lifespan extension.

### Post-reproductive BEC-1 suppression mediates longevity through its canonical role in autophagy

BEC-1 exists in at least two distinct protein complexes with either CED-9/Bcl-2 or VPS-34 and EPG-8, respectively inhibiting caspase-mediated apoptosis or promoting autophagic nucleation (Takacs-Vellai et al. 2005; Yang and Zhang 2011). Inactivation of *bec-1* triggers CED-4 dependent apoptotic cell death (Takacs-Vellai et al. 2005). Combinatorial suppression of *bec-1* and *ced-4* at day 9 did not alter *bec-1* meditated MTL extension (Fig. 3A, and Supplemental Fig. S3C), suggesting an apoptosis independent longevity mechanism. However, RNAi against either *epg-8* or *vps-34* at day 9 caused lifespan extensions similar to *bec-1* knockdown (Fig. 3B). Inactivation of *epg-8* or *vps-34* from the first day of adulthood caused a significantly reduced lifespan (Supplemental Fig. S3A), implying that AP is a common feature of the VPS-34/BEC-1/EPG-8 autophagic nucleation complex. We then investigated mitophagy as another principle form of autophagy involving BEC-1 (Palikaras et al. 2015), by inactivating either *dct-1* or *pink-1* at day 9. Neither condition effected MTL, indicating that mitophagy is unlikely to be involved (Supplemental Fig. S3B). Late-life reduction in autophagy via *bec-1* may mediate its longevity effects through upregulation of the ubiquitin-proteasome system (UPS), the other major protein degradation pathway. However, day 9 inactivation of *bec-1* neither impacted the chymotrypsin-like activity of the proteasome nor the abundance of lysine 48-linked polyubiquitin chains that target proteins for degradation (Supplemental Fig. S3, D and E). Next, we examined whether day 9 inactivation of *pha-4* or *bec-1* causes unfolded protein or oxidative stress responses. Expression levels of genes involved in both stress resistance pathways were not significantly altered, which argues against their involvement in the observed longevity phenotype (Supplemental Fig. S3, F and G).

**Fig. 3.**
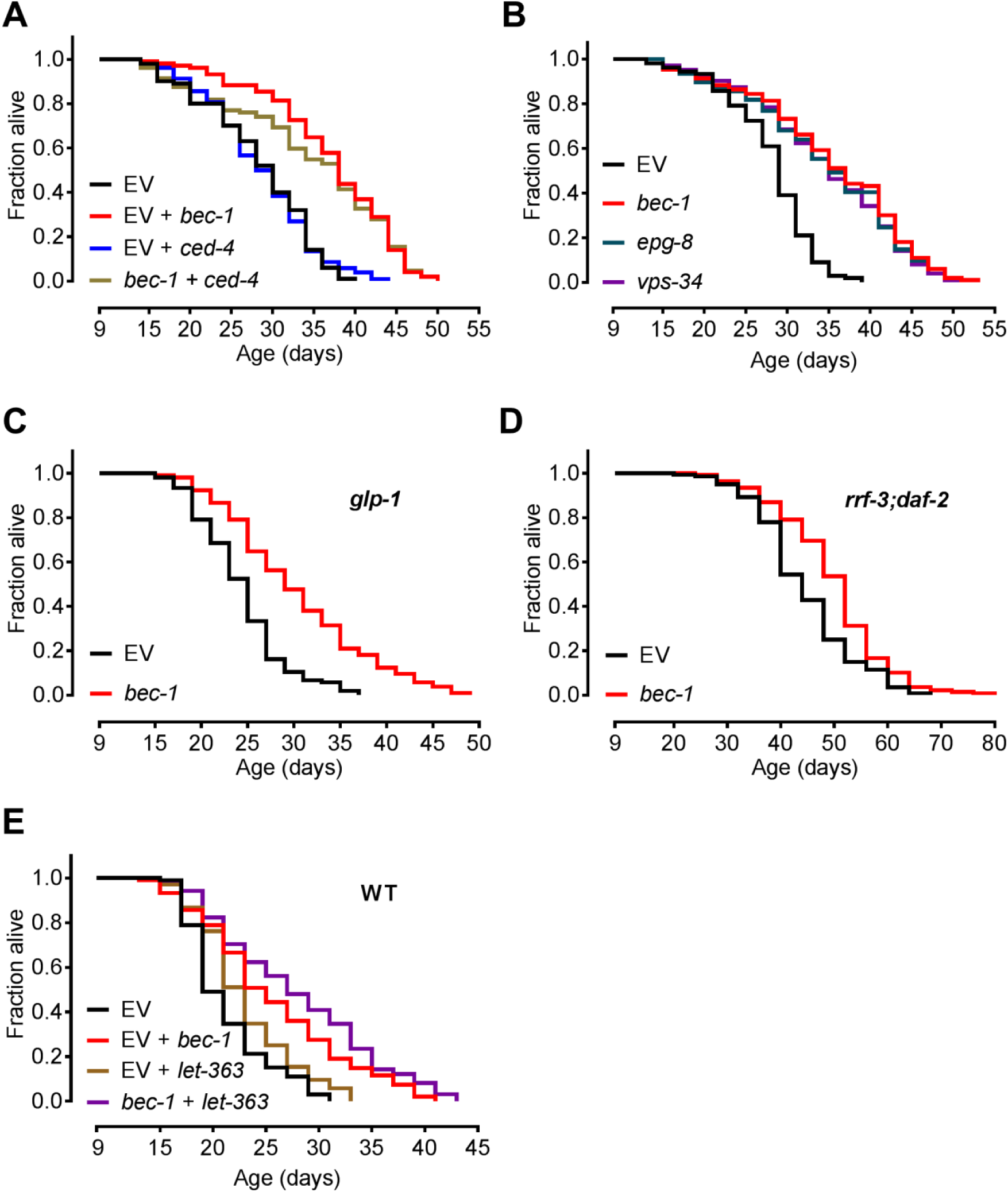
Inactivation of *bec-1* extends lifespan through autophagy and is independent of classical longevity pathways. (A) *bec-1* RNAi treatment shows no change in effect on MTL with simultaneous *ced-4* RNAi treatment at day 9 in *rrf-3(pk1426)* worms. (B) *bec-1*, *vps-34* and *epg-8* equally extend the MTL of *rrf-3(pk1426)* worms when inactivated at day 9. (C) Day 9 RNAi against *bec-1* extends the MTL of *glp-1(e2141)* worms. (D) Day 9 RNAi against *bec-1* extends the MTL of *rrf-3(pk1426)*; *daf-2(e1370)* worms. (E) Combined inactivation of *bec-1* and *let-363*(TOR) at day 9 in WT(N2) worms further extends their MTL compared to *let-363* inactivation alone. — Days represent days post first egg lay. Lifespan statistics, Supplemental Table 6.

### Post-reproductive bec-1 inactivation further extends lifespan in autophagy dependent longevity mutants

We tested if *bec-1* inhibition still exhibits an AP phenotype in worms with reduced germline-, insulin-or TOR-signaling. Longevity mediated via these pathways has been shown to depend on increased autophagy (Sheaffer et al. 2008; Lapierre et al. 2011; Meléndez et al. 2003; Lapierre et al. 2013; Hars et al. 2007; Hansen et al. 2008). In contrast to observations in young worms (Lapierre et al. 2011; Hansen et al. 2008), the germline-less *glp-1* and insulin/IGF-1-like receptor *daf-2* mutants showed a significant MTL extension upon day 9 *bec-1* RNAi (Fig. 3, C and D). Additionally, *bec-1* inactivation in *daf-2;daf-16* double mutants extended MTL (Supplemental Fig. S3H). Combinatorial inactivation of TOR/*let-363* and *bec-1* at day 9 also showed an additive effect on MTL in WT worms (Fig. 3E). Similar to reports in young animals (Mizunuma et al. 2014), day 9 RNAi against *let-363* in *rrf-3* mutants did not significantly extend MTL, but also did not impact *bec-1* mediated longevity (Supplemental Fig. S3, C and I). Our data shows a converse modulation of lifespan by *bec-1*, even in mutants whose longevity normally strictly depends on autophagy.

### Inhibition of the autophagic cascade up until vesicle nucleation extends lifespan

After autophagy induction via the serine/threonine protein kinase UNC-51 and vesicle nucleation via the BEC-1 complex, autophagy is broken down into further steps that include vesicle expansion, lysosomal fusion and degradation. We inactivated representative genes from each step at day 9 and examined lifespan effects (Fig. 4A, Supplemental Table 3, and Supplemental Fig. S4, A to E). Post-reproductive inhibition of autophagy genes caused a strong MTL extension up to and including the step of vesicle nucleation. After this, inhibition of the later autophagic flux did not similarly increase lifespan. Interestingly, day 9 inhibition of *lgg-1*, but not its homologue *lgg-2* (Alberti et al. 2010), significantly reduced lifespan (Supplemental Fig. S4C). Shortly after *lgg-1* inactivation, we further detected a sick and sluggish phenotype in young as well as in old worms (data not shown). This negative phenotype was not observed upon post-reproductive inactivation of any other tested autophagy gene. Notably, this included genes that are critical for the functionality of LGG-1 in autophagy, such as ATG-4.1 (Wu et al. 2012), ATG-7 (Kim et al. 1999), and LGG-3 (Mizushima et al. 1998) (Supplemental Fig. S4C). Also, *lgg-1* day 9 RNAi still exerted negative effects when autophagy was independently repressed through simultaneous *bec-1* inactivation (Supplemental Fig. S4, F and G). Combined, these results suggest critical non-autophagy functions of LGG-1 and that vesicle nucleation is integrally linked to the longevity phenotype.

**Fig. 4.**
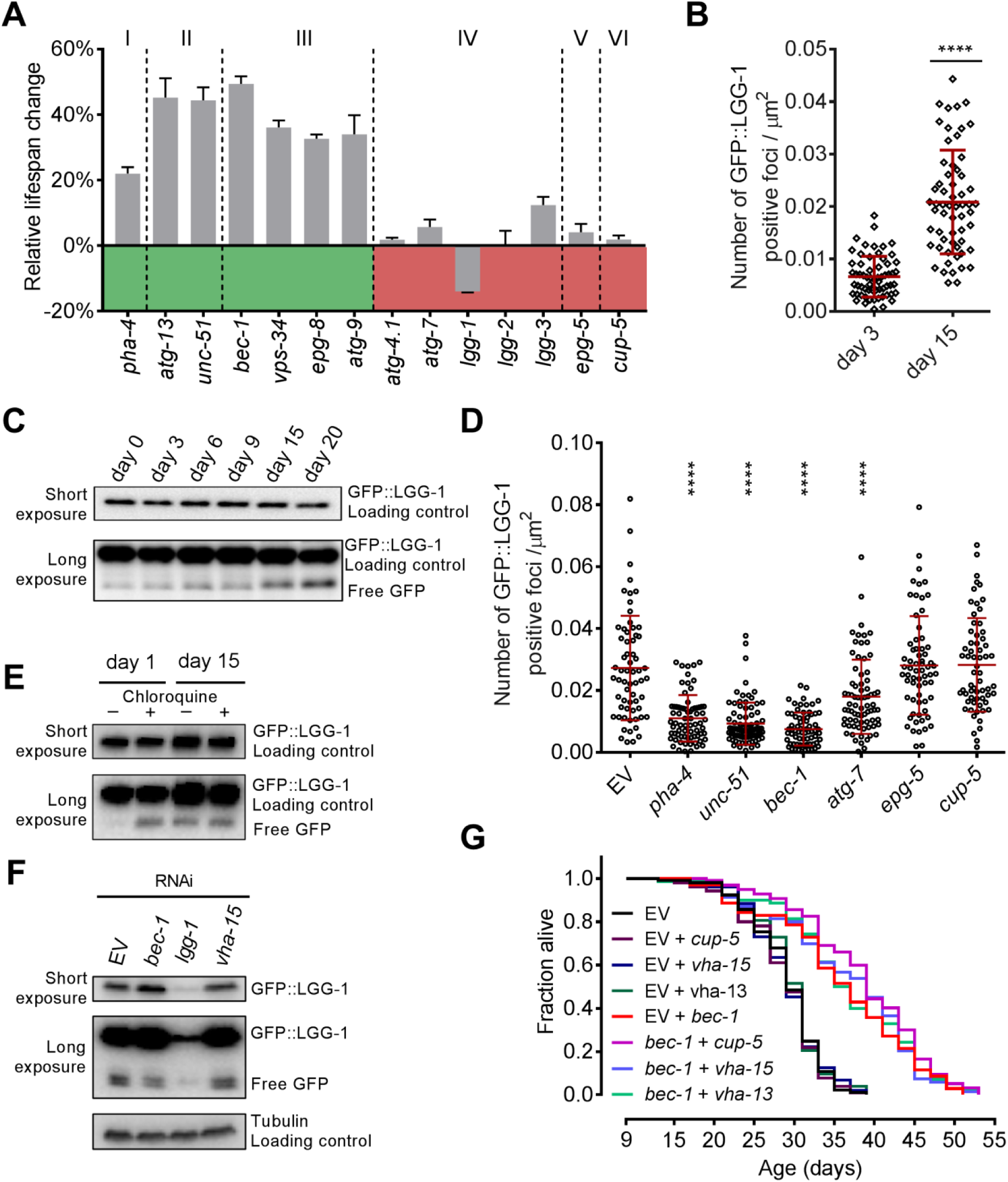
Autophagy is dysfunctional in aged worms and inhibition of its initiation increases lifespan. (A) Day 9 RNAi against genes involved in autophagosome: regulation (I), induction (II), nucleation (III), expansion/maturation (IV), lysosomal fusion (V), and degradation (VI). Showing relative percentage change in MTL +/-SEM compared to control. Statistics, Supplemental Table 3. Gene knockdown validation (Supplemental Fig. S4E). (B) Quantification of GFP::LGG-1 foci over time in hypodermal cells of *rrf-3(pk1426); lgg-1p::GFP::lgg-1*. Foci were quantified in young and old worms from images at 100x magnification (N=50). Red lines show median and interquartile range (**** = p<0.0001). (C) Representative western blot of free GFP across life balanced to total GFP::LGG-1. The cleaved GFP forms correspond to the GFP::LGG-1 degradation products in the autolysosome. (D) Day 15 Quantification of GFP::LGG-1 positive foci in hypodermal cells of *rrf-3(pk1426); lgg-1p::GFP::lgg-1* worms following day 9 RNAi knockdown. Foci were quantified from images with 100x magnification (N=50). Red lines show median and interquartile range (**** = p<0.0001, *** = p<0.001, ** = p<0.01). LGG-1 knockdown validation (Supplemental Fig. S4G). (E) Representative western blot of free GFP, balanced to GFP::LGG-1, in young and old worms treated with chloroquine. The cleaved GFP forms correspond to the GFP::LGG-1 degradation products in the autolysosome (F) Representative western blot of day 9 RNAi against: *bec-1, vha-15* and *lgg-1* and its effect on free GFP compared to GFP:LGG-1 measured at day 15. The cleaved GFP forms correspond to the GFP::LGG-1 degradation products in the autolysosome (G) Day 9 inactivation of lysosomal degradation or acidification in *rrf-3(pk1426)* has no effect on MTL, or on the MTL increase mediated by *bec-1* inhibition when used in combination. — All western blots used antibodies against GFP. Days represent days post first egg lay. Lifespan statistics, tables S8 and S9.

### Aged worms exhibit a dysfunctional late autophagic flux

Upon autophagosome formation, GFP::LGG-1 changes its diffuse cytoplasmatic localization to distinct foci that represent autophagic structures (Meléndez et al. 2003; Kang et al. 2007). Interestingly, we detected an age-dependent increase in these autophagic structures by quantifying GFP::LGG-1 foci in the hypodermis (Fig. 4B and Supplemental Fig. S5A). Next, we monitored the two autophagy reporters GFP::LGG-1 and BEC-1::RFP across age by western blotting. We observed a strong depletion of GFP::LGG-1 and BEC-1::RFP protein pools in old worms (Supplemental Fig. S5, B and C), which was not reflected in their endogenous mRNA levels (Supplemental Fig. S5D). After fusion of the autophagosome with the lysosome, Atg8/LC3/LGG-1 reporter proteins are degraded by lysosomal hydrolases (Klionsky et al. 2016). Conversely, the coupled GFP fragment accumulates in the vacuole as it is relatively resistant to the vacuolar enzymes (Klionsky et al. 2016). The efficiency of lysosomal degradation determines how long the free GFP moiety can be detected. Thus, the ratio of full-length protein to its cleaved reporter indicates the status of the flux (Klionsky et al. 2016). Compared to the full-length GFP::LGG-1 pool, free GFP increased from day 0 to 20 of adulthood (Fig. 4C). We obtained similar results with worms expressing the autophagic reporter protein BEC-1::RFP (Supplemental Fig. S5C).

Together, our data supports three alternative autophagic flux states in old worms: an increased formation of autophagosomes due to an enhanced autophagy induction, inefficient completion of autophagy resulting in accumulation of autophagic structures, or a combination of both (Klionsky et al. 2016). To discriminate between these options, we monitored GFP::LGG-1 foci levels present in old worms upon day 9 inhibition of proteins involved at different stages of the autophagic flux. Inactivation of *pha-4*, *unc-51, bec-1*, and *atg-7* at day 9 significantly reduced autophagosome numbers (Fig. 4D and Supplemental Fig. S6). As *epg-5* and *cup-5* function in lysosomal fusion and degradation respectively, their inactivation is associated with an accumulation of GFP::LGG-1 foci (Tian et al. 2010; Sun et al. 2011). In contrast, we did not observe a significant increase in GFP::LGG-1 foci upon *epg-5* or *cup-5* RNAi from day 9. This lack of accumulation suggests a blocked late autophagic flux leading to inefficient autophagy completion. Next, we simulated an inefficient completion of autophagy through genetic or pharmacological perturbation of autophagic degradation by neutralizing lysosomal pH in young and old worms. In young animals, chloroquine treatment or the inactivation of the vacuolar ATPase subunit VHA-15 caused a significant increase in free GFP (Fig. 4E and Supplemental Fig. S7A). However, these treatments did not increase free GFP levels in aged worms (Fig. 4, E and F). Hence, the level of free GFP in young worms with an artificially perturbed late autophagic flux mimics the intrinsic status in old worms.

To further substantiate our analysis of the autophagic flux across aging, we monitored the autophagic substrate SQST-1/p62. Blocking autophagy leads to the accumulation of the adaptor protein SQST-1, which recognizes and loads cargo into the autophagosome for degradation (Tian et al. 2010). We observed a marked increase in SQST-1 with age (Supplemental Fig. S7B), which is supported by a recent proteome analysis (Narayan et al. 2016). We did not detect a matched increase in *sqst-1* mRNA expression levels (Supplemental Fig. S7C), which speaks for an accumulation on the protein level. RNAi against different autophagy genes at day 9 did not cause a marked increase in SQST-1 (Fig. S7D). Yet, we observed a striking SQST-1 accumulation upon *bec-1* inactivation at day 0 (Supplemental Fig. S7E). The progressive accumulation of SQST-1::GFP (Supplemental Fig. S7B) in spite of increasing numbers of autophagic structures (Fig. 4D) in aged worms, corroborates a dysfunction of late-life autophagy. Transcriptional upregulation of autophagy genes could indicate increased autophagic induction. Using a sequencing approach, we did not observe a systematic upregulation of autophagy genes over age, in line with proteomic observations (Walther et al. 2015). Notably, we detected a ∼20% increase in *bec-1* expression across aging (Supplemental Fig. S5D).

Our findings imply two possible mechanisms by which post-reproductive inhibition of autophagic vesicle nucleation could be beneficial: either inhibition alleviates the pressure on the autophagic system to restore productive autophagy or it prevents detrimental effects from the early termination of dysfunctional autophagy processes. Hence, we tested for restored productive autophagy by investigating if *bec-1* mediated lifespan extension requires lysosomal degradation. Simultaneous inactivation of *bec-1* with either *cup-5*, *vha-13* or *vha-15* at day 9 did not affect the *bec-1* longevity phenotype (Fig. 4G). While we could successfully inhibit lysosomal acidification in old worms, inactivation of *vha-13* or *vha-15* at day 9 had no effect on lifespan (Supplemental Fig. S7F and S4D). Conversely the same inactivation strongly reduced the lifespan of young worms (Supplemental Fig. S7G). Thus, *bec-1* inactivation at day 9 does not restore productive autophagy. Its AP effects most likely arise as harmful consequences of impaired late stage autophagy.

### Aged WT worms show reduced neuronal RNAi and decreased longevity gains upon inhibition of autophagic nucleation

We also examined the effect of inhibition of autophagosome nucleation in WT worms. In agreement with published data (Sheaffer et al. 2008; Meléndez et al. 2003), RNAi against *bec-1* or *pha-4* results in a significant reduction of lifespan early in life (Supplemental Fig. S8, A and B). Conversely, day 9 inhibition of autophagy genes in WT resulted in similar lifespan extending trends as in the *rrf-3* mutant but with a weaker strength (Supplemental Fig. S8C). *bec-1, vps-34, unc-51* and *atg-9* all extended lifespan approximately by 30%. Notably, we did not observe a lifespan extension following *pha-4* inhibition from day 9 (Supplemental Fig. S8C). However, as observed in *rrf-3* mutants, combined inactivation of *pha-4* and *bec-1* in WT resulted in reduced longevity compared to *bec-1* RNAi alone (Supplemental Fig. S8D), implying a functional but weak effect of *pha-4* inhibition. Combined, this suggests that the weaker effects on longevity in old worms compared to young worms could derive from differences in RNAi efficacy between WT and *rrf-3* mutant worms. Loss of functional RRF-3 enhances global sensitivity to RNAi in young worms (Simmer et al. 2002). Upon day 9 RNAi against *pha-4* and *bec-1*, we likewise observed a reduced knockdown efficiency in WT compared to *rrf-3* mutants at day 12, which equalized by day 15 (Supplemental Fig. S8E). Moreover, RNAi is particularly effective in the neurons of *rrf-3* mutants compared to WT, which are largely refractory to neuronal RNAi (Simmer et al. 2002). We confirmed that this enhanced neuronal RNAi sensitivity persists in aged *rrf-3* worms, using *unc-119::gfp* as a neuronal reporter (Supplemental Fig. S8F). Combined with the striking MTL differences between the two strains, we were prompted to investigate neuronal contribution to the observed longevity.

### Inhibition of bec-1 extends healthspan and lifespan through neurons

We performed lifespan experiments in the *sid-1(-); unc-119p::sid-1(+)* mutant strain that is hypersensitive to RNAi in neurons but refractory in all other tissues (Calixto et al. 2010). Intriguingly, neuron-specific inactivation of *bec-1* and *vps-34* at day 9 strongly extended MTL by up to 57% (Fig. 5A). *atg-7* did not alter lifespan, and *lgg-1* significantly reduced MTL (Fig. 5A). We observed a significant reduction in lifespan upon neuron-specific *bec-1* RNAi at day 0 (Supplemental Fig. S9A), which indicates a conserved neuronal AP effect for *bec-1*. These lifespan effects mirror those seen in our *rrf-3* mutant experiments. Notably, inactivation of *bec-1* at day 9, specifically in the germline niche, gut, muscle or hypodermis, did not affect lifespan (Supplemental Fig. S9B). These results suggest that enhanced neuronal RNAi is likely the main contributor to extended longevity in *rrf-3* mutants. Taken together, our observations imply that inhibition of autophagic nucleation extends lifespan primarily through neurons.

**Fig. 5.**
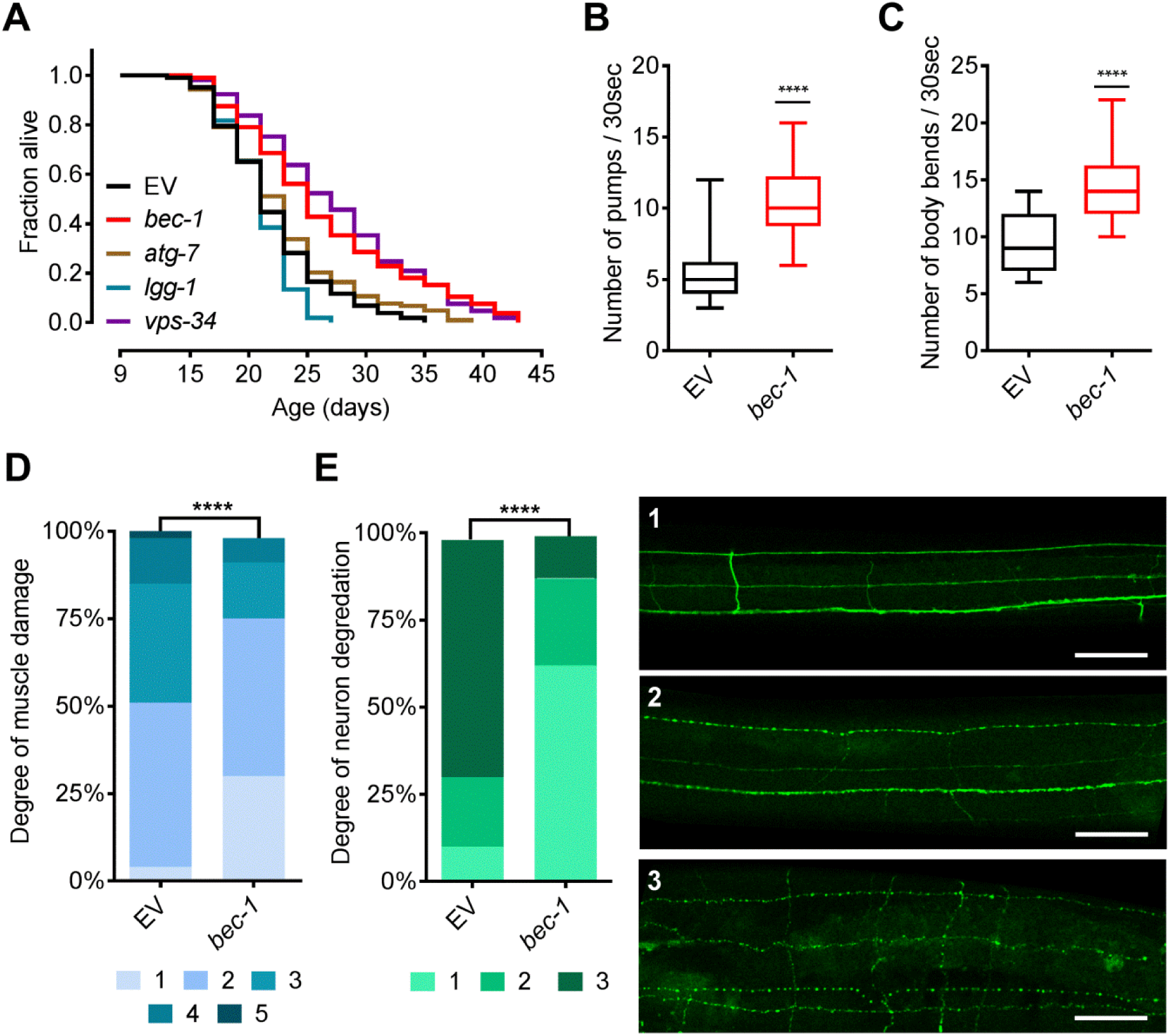
Neuronal inhibition of *bec-1* mediates health and lifespan effects. (A) Day 9 RNAi against members of the autophagic flux extend MTL in *sid-1*(pk3321); *unc-119p::sid-1* worms. (B) Day 20 quantification of phalloidin stained muscle damage following day 9 RNAi against *bec-1* in *sid-1*(pk3321); *unc-119p::sid-1* worms. Scoring system is the same as used in Fig. 1F (N=100) (**** = p<0.0001). (C) Quantification of worm mobility measured by body bend counts in *sid-1*(pk3321); *unc-119p::sid-1* worms at day 20. Data depicted as a box plot with min to max values. RNAi against *bec-1* was initiated at day 9. (N=50) (**** = p<0.0001). (D) Day 20 quantification of pharynx pumping rate in *sid-1*(pk3321); *unc-119p::sid-1* worms. Data depicted as a box plot with min to max values. RNAi against *bec-1* initiated at day 9. (N=50) (**** = p<0.0001). (E) Day 20 quantification of longitudinal neuron morphology and integrity in *unc-119p::GFP; rrf-3(pk1426)* worms treated with RNAi against *bec-1* at day 9. Neurons scored on a three-point scale based on the amount of visible bubbling and neuron integrity. Neurons quantified from images at 40x magnification (N=100). Scale bar 40 μm. (**** = p<0.0001). — Days represent days post first egg lay. Lifespan statistics, Supplemental Table 11.

**Fig. 6.**
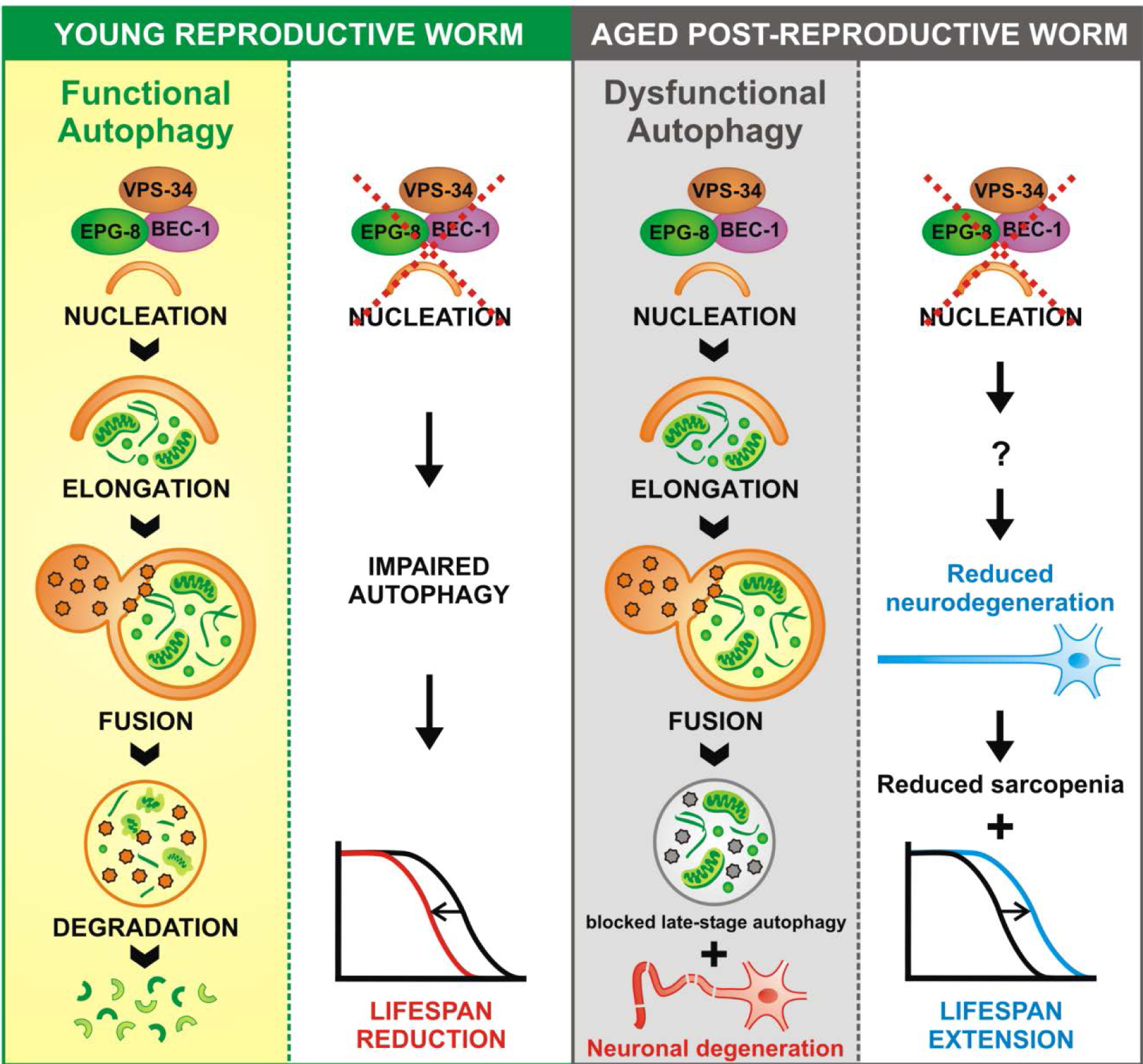
Model depicting age associated antagonistic effects of autophagy on longevity. (Left Panel) Young worms have functional autophagy and inhibition of vesicle nucleation in these young worms shortens lifespan. (Right Panel) Old worms still have effective autophagy induction but exhibit an age related dysfunction in autophagic degradation. This is accompanied by age-associated neurodegeneration. Inhibition of autophagic nucleation in old worms, through an unknown mechanism, leads to improved neuronal integrity and an accompanying decrease in sarcopenia. This improvement in health acts either causally or in concert with an increase in post-reproductive lifespan.

Next, we examined if neuron-specific inactivation of the nucleation complex via *bec-1* accounts for the improvement in global health. Day 9 inactivation of *bec-1* in the neurons promoted a youthful muscle structure, improved mobility, and an increased pumping rate (Fig. 5, B to D). However, the same inactivation in solely muscle tissue failed to generate similar improvements (Supplemental Fig. S9, C to E). Additionally, we examined if inhibition of the early flux through *bec-1* would elicit changes in neuronal integrity. Intriguingly, *bec-1* inhibition at day 9 resulted in more youthful neurons with a significant reduction in axonal blebbing and fragmentation (Fig. 5E and Supplemental Fig. S9F). Overall, our data suggests that neuroprotective effects of *bec-1* inactivation mediate an improved muscle function and integrity.

## Discussion

### Novel screening approach for AP in post-reproductive C. elegans

Prevalent age-associated disorders such as cardiovascular diseases, cancer, and neurodegenerative diseases have a strong genetic contribution (Kulminski et al. 2016). With the increase in our aging population, disease prevention as opposed to disease treatment of the elderly is a promising approach for future medicine. However, such preventive therapies require an improved understanding of the genetics of aging. Based on the AP theory of aging, our study illustrates that genetic observations made in young organisms cannot easily be translated into an aged context. In contrast, here we show that key autophagy genes required for normal lifespan in young worms antagonistically shorten life in post-reproductive worms. In line with this finding, recent human genome-wide association studies showed that disease risk associated with a given allele is highly sensitive to chronological age (Kulminski et al. 2016; Rodríguez et al. 2017). Yet, high-throughput screens for genes affecting lifespan in *C. elegans* largely ignored the impact of chronological age, as they were carried out in young worms (Ni and Lee 2010; Klass 1983). Our study is the first to report a post-reproductive RNAi screen for genes that behave according to the AP theory of aging. This novel approach led to the identification of 30 previously unknown longevity genes that increase lifespan upon post-reproductive inactivation. These findings indicate that AP is a common genetic mechanism that modulates health and lifespan in an age-dependent manner.

### Dysfunctional autophagy in aged worms

Our data provides evidence for active autophagic induction in old worms, resulting in accumulation of autophagic foci that are not cleared likely due to a blockage in the late flux. This is in line with observations in aging mice (Terman 1995; Del Roso et al. 2003). A potential blockage of the flux is further evidenced by the absence of an increase in autophagosome structures or free GFP upon day 9 inactivation of the autophagic degradation step (Figure 4, E and F). The accumulation of the autophagy receptor SQST-1 in old worms harkens back to studies that showed widespread protein accumulation in aged *C. elegans* (Walther et al. 2015; David et al. 2010), and implicates ineffective late-life autophagy. We also observed that genes crucially involved in autophagy degradation are dispensable late in life and have no effects on longevity (Supplemental Fig. S4D), which further argues for a dysfunctional autophagic process in aged worms. Moreover, the significant reduction of autophagic structures upon day 9 inactivation of the early autophagic flux (Fig. 4D and Supplemental Fig. S6) indicates continuing vesicle nucleation in aged worms. Further, the age-associated increase in *bec-1* mRNA (Supplemental Fig. S5D), while BEC-1 and LGG-1 protein decreases (Supplemental Fig. S5, B and C), could suggest an elevated autophagosome formation with age. Altogether, our findings indicate active and possibly elevated vesicle nucleation combined with a blockage downstream of autophagosome biogenesis in aged worms.

Autophagy represents a quality control system that targets misfolded proteins and thereby prevents their toxic intracellular accumulation. Inhibition of the system in mice leads to the accumulation of toxic protein aggregates and cellular degeneration of cardiomyocytes and neurons (Nakai et al. 2007; Komatsu et al. 2006). As such, autophagy is a major factor that delays aging (Rubinsztein et al. 2011). During aging organisms are increasingly challenged with large protein aggregates that are poor substrates to the proteasome (Walther et al. 2015; David et al. 2010). Thus, one would expect that autophagic activity increases with age. Yet, many studies detected decreasing autophagic activity during aging (Rubinsztein et al. 2011). It is still unclear how worms could compensate for a dysfunctional autophagic recycle machinery in old age. Proteomic data across lifespan shows an increasing abundance of proteasome components with age (Walther et al. 2015), suggesting that an upregulation of the UPS could be one possibility by which aged animals maintain proteostasis. However, we did not detect further upregulation of the UPS activity upon *bec-1* inactivation from day 9 (Supplemental Fig. S3, D and E), which may suggests that the UPS is already working at its maximal capacity in aged worms. Hence, dysfunctional autophagy in late-life may activate compensatory mechanisms such as increased UPS and increased protein sequestration mechanisms (Walther et al. 2015) to improve overall proteostasis, allowing the organism to survive.

### Autophagosome nucleation exhibits AP

Through downstream analysis of our screening hit *pha-4*, we identify that the observed longevity effects are mediated through inhibition of the key autophagy gene *bec-1*. Next, we found that BEC-1 mediates its longevity effects through the VPS-34/BEC-1/EPG-8 autophagic nucleation complex rather than through its interaction with the antiapoptotic protein CED-9/Bcl-2. Notably, BEC-1 and VPS-34 have also been shown to function together in endocytosis (Ruck et al. 2011). However, EPG-8 is not involved in this process (Yang and Zhang 2011), arguing against endocytic involvement. Hence, our data suggests that post-reproductive inactivation of *bec-1* mediated its longevity through its role in autophagy. Surprisingly, day 9 inactivation of *bec-1* also extends lifespan in long-lived worms with reduced germline-, insulin-or TOR-signaling, whose longevity was previously linked to enhanced autophagy (Sheaffer et al. 2008; Lapierre et al. 2011; Hansen et al. 2008; Meléndez et al. 2003; Lapierre et al. 2013; Hars et al. 2007). Our findings do not contradict these reports, as they studied longevity effects of autophagy in young worms. Rather, our data suggests that even in the presence of lifespan extending mutations, autophagy eventually becomes dysfunctional at old age, altering the process from beneficial to harmful.

Autophagy genes involved in steps subsequent to vesicle nucleation show minor or no longevity effects upon day 9 inactivation, even if their inhibition also reduces autophagosome numbers. This is illustrated by ATG-7, which acts immediately downstream of vesicle nucleation and is needed for autophagic vesicle elongation (Kim et al. 1999). Similar to results in young worms (Kang et al. 2007), we observed significantly reduced GFP::LGG-1 foci numbers upon day 9 inactivation of ATG-7 (Fig. 4D). However, day 9 RNAi against *atg-7* had only minor effects on lifespan. Similarly, day 9 inactivation of LGG-1, which by itself is critical for vesicle expansion and completion (Meléndez et al. 2003), did not positively affect *C. elegans* lifespan. Interestingly, *lgg-1* is the only autophagy gene tested that shows negative effects post day 9. This is potentially due to roles of LC3/LGG-1 outside autophagy, such as in the ER-associated degradation (ERAD) tuning process (De Haan et al. 2010). As such, there seems to be a decoupling of autophagosome formation and longevity. Our data indicates that autophagosome numbers are presumably not critical for lifespan extension but rather the stage at which the flux is inhibited. This suggests that, in the context of age-related autophagic dysfunction, the vesicle nucleation event causes harmful lifespan shortening effects.

Notably, post-reproductive inactivation of *pha-4* resulted in weaker lifespan extension than observed for *bec-1* RNAi and also reduced *bec-1* longevity upon combinatorial inactivation from day 9 (Fig. 2, A and B, Supplemental Fig. S8, C and D). Besides its role as an upstream transcriptional regulator of *bec-1*, the transcription factor PHA-4 also targets gene classes that are involved in metabolic processes and defense responses (Zhong et al. 2010). In addition to reducing autophagy, one would expect that post-reproductive inactivation of PHA-4 results in a dysregulation of its non-autophagy target genes. This likely explains the reduced longevity of *pha-4* compared to *bec-1* RNAi, as well as its subtractive lifespan effect when co-repressed with *bec-1*. Similarly, reduced positive effects through autophagy inhibition in WT worms are probably canceled out by negative effects of PHA-4 inactivation, resulting in no net effect on WT lifespan upon day 9 *pha-4* RNAi.

### Inhibition of the autophagy nucleation complex extends lifespan through neurons

Dysfunctional autophagy has been linked to neuronal degeneration in cell culture as well as in animal models of neurodegenerative diseases (Nixon 2013). Similarly, accumulation of dysfunctional autophagic structures were observed in human Alzheimer’s, Parkinson’s, and Huntington’s disease patients (Nixon 2013). Interestingly, the late autophagic flux rather than the process of autophagosome formation has been frequently reported to be impaired in these neurodegenerative diseases. However, the exact mechanistic link between impairment of the late autophagic flux and neurodegeneration is still a matter of debate and seems to largely depend on the disease context (Nixon 2013). Proposed mechanisms that link late flux dysfunction to neuronal damage include: defective retrograde transport, harmful accumulation of autophagosomes and autolysosomes, ineffective organelle clearance, increased ROS production, incomplete degradation as a source of cytotoxic products, and cell death through leakage of lysosomal enzymes (Nixon 2013). Interestingly, the inhibition of autophagosome formation has been shown to ameliorate neurodegeneration under disease conditions that show a pathologic accumulation of autophagic vesicles (Lee and Gao 2009; Yang et al. 2007). We demonstrate that neuron-specific inactivation of autophagic flux genes recapitulated the longevity effects seen through their global inactivation, implying that autophagy in the nervous system is equally impaired as in the whole organism. Further research is still needed to establish the specific autophagy status in aged *C. elegans* neurons. Our study suggests that age-associated dysfunctional autophagy causes neurodegeneration and shortens organismal lifespan. Here we provide the first evidence that inhibition of autophagic vesicle nucleation ameliorates neuronal decline during aging. Further, we show that neuron-specific inactivation of the vesicle nucleation complex through *bec-1* is sufficient to extend health-and lifespan in aging *C. elegans*. In contrast to other studies that show reduced neuronal decline upon inactivation of genes involved in autophagic vesicle elongation (Lee and Gao 2009; Yang et al. 2007), our neuroprotective and longevity phenotype is exclusively linked to genes that induce or participate in the process of vesicle nucleation.

An age-related decline in proteostasis has been associated with the accumulation of misfolded and aggregated proteins that are widely implicated in neurodegeneration (Nixon 2013). When autophagy is dysfunctional in old age, neurons may largely depend on protein sequestration or exophere formation to reduce cytotoxic effects (Walther et al. 2015; Melentijevic et al. 2017). On a mechanistic front, ongoing vesicle nucleation feeding into a dysfunctional autophagic flux may interfere with these neuroprotective pathways (Melentijevic et al. 2017; Arrasate et al. 2004). Along these lines, we show that post-reproductive inactivation of *bec-1* preserves neuronal integrity. However, the potential involvement of these sequestration pathways in our phenotype remains to be investigated. Also, we cannot exclude unknown lifespan-shortening signaling events emanating from the autophagic vesicle nucleation complex in late-life.

Our study shows a global blockage of autophagy at the later steps of autophagosomal degradation in aged worms. Inhibiting this process at the critical step of vesicle nucleation through the BEC-1/VPS-34/EPG-8 complex alleviates harmful consequences of the age-related autophagic dysfunction. We show that these beneficial effects are mediated through neurons and result in reduced neurodegeneration, reduced sarcopenia and increased lifespan (Fig 6).

## Materials and Methods

### C. elegans strains

*C. elegans* strains were maintained at 20°C using standard procedures (Brenner 1974) unless indicted differently. A complete list of the strains used in this study is in the Supplemental Material.

### RNAi screen

800 dsRNA expressing HT115 *E. coli* bacteria specific to *C. elegans* gene regulatory factors was prepared from the Ahringer and Vidal libraries (Supplemental Table 1). All clones were sequence validated using the M13 forward primer. Library was grown overnight in 2x YT media in 96 deep well plates and seeded onto 24 well NGM agar plates with 100 μg/ml Ampicillin and 1 mM β-D-isothiogalactopyranoside (IPTG) at 2x density. Minimal 4 replicate wells per RNAi clone were seeded. Non-targeting controls comprised of 4 plates of both EV and *gfp* RNAi. *gfp* RNAi was as kind gift from Scott Kennedy. *rrf-3* mutant worms were synchronized via liquid culture until day 9, cleaned and sorted with the COPAS Biosorter with 20 worms/well. Live/dead scoring of all wells was performed on day 32, when most control animals were dead. Scoring was performed manually by flushing each well with M9 buffer and immediately counting moving and dead worms. Only worms which could be identified as live or dead were scored, missing worms were not scored. Contaminated wells or offspring containing wells were not scored, combined these conditions represented ∼10% of all wells.

### Lifespan assays

Worms were synchronized using liquid culture sedimentation. dSRNA expressing bacteria were grown overnight, seeded on NGM agar plates with 100 μg/ml Ampicillin and 1mM IPTG, and incubated at RT over night. Day 0 was determined with appearance of first internal eggs. A total of 105 -140 animals were placed on 3 to 4 replicate plates with 35 worms per 6 cm plate. Worms were picked to new plates and scored every two days. For double RNAi treatment, bacteria were diluted in a 1:1 ratio before being plated at 1x density. All assays were performed at 20 °C. For temperature sensitive *glp-1* mutant, worms were maintained restrictive temperature of 25 °C until day 0, when transferred to 20 °C. Lifespan assays indicated in supplementary table as “plate only”, worms were maintained and synchronized only on plates. Worms were scored as alive until there was no movement after repeated prodding. Worms were censored when: crawled off plate, bagging, bursting, dropping on transfer, contamination, and burrowing. All RNAi clones used for lifespan assays are from the Ahringer libary and were sequence validated using the M13 forward primer.

### LGG-1::GFP microscopy

Worms were grown as described and transferred to respective RNAi treatments at day 9. On the day of analysis, 40 worms from each knockdown condition were paralyzed using 0.5% NaAz and mounted on a 2% agar pad on a glass microscope slide. GFP signals were acquired using a STED super-resolution microscope (Leica) at 100x magnifications. Images were taken of the hypodermis which was identified by locating the plane between the muscle and cuticle. At least 60 total regions were imaged from 50 different worms per replicate with two biological replicates. All images used for comparative quantifications were taken on the same day, with the same settings and by the same user.

### Pharynx imaging and analysis

Worms were grown as described and transferred to respective RNAi treatments at day 9. On day 20, 50 aged-synchronized worms from each knockdown condition were paralyzed using 0.5% NaAz and mounted on a 2% agar pad on a glass microscope slide. The images of the pharynges were acquired on a SP5 microscope (Leica) using the Differential Intense Contrast (DIC) filter and a 63x/1.4 NA Oil immersion objective. At least 30 whole worms per condition were imaged in each replicate with three technical replicates. Images were scored blind for pharynx degradation based on a three-point scale of damage related to changes in the structure of the corpus, isthmus or terminal bulb. Worms with no obvious damage to any of the three parts were scored as 1, worms with damage to only one or two parts were scored 2, and worms with damage to all three parts were scored 3. For comparative analysis all worms were imaged at the same time, with the same settings and by the same user. This experiment was carried out in two independent biological replicates.

### Muscle cell imaging and analysis

Worms were grown as described and transferred to respective RNAi treatments at day 9. On day 20, 30 aged-synchronized worms from each knockdown condition were incubated in fixation buffer (160 mM KCl, 100 mM Tris HCl pH 7.4, 40 mM NaCl, 20 mM Na_2_EGTA, 1mM EDTA, 10 mM Spermidine HCl, 30 mM Pipes pH 7.4, 1% Triton-x-100, 50% Methanol) at room temperature for 1 hour rotating. Worms were washed twice with PBS and incubated in a 1:200 dilution of Phalloidin–Atto 565 (Sigma) in PBS-0.5% Triton-x-100 for 4 hours at room temperature rotating. Worms were then mounted on a 2% agar pad on a glass microscope slide. The 561 nm signals were acquired using a STED CW super-resolution microscope (Leica) and a 63x/1.4 NA Oil immersion objective. At least 40 images, comprising 2-4 cells, from 20 different worms were imaged per replicate with three technical replicates. Images were scored blind based on a five-point scale of muscle fiber degradation. Cells showing no obvious degradation were scored as 1, cells with kinks or striations were scored as 2, cells with small lesions combined with striations or other damage were scored 3, cells with muscle fiber breaks, gross striations and lesions were scored as 4, and cells where the muscle fibers were no longer intact were scored as 5. For comparative analysis, all worms were imaged at the same time, with the same settings and by the same user. This experiment was carried out in two independent biological replicates.

### Pharynx pumping assay

Worms were grown in liquid culture and transferred to respective RNAi treatments at day 9. On day 20, 50 aged-synchronized worms from each knockdown condition were transferred individually on an agar plate seeded with a bacterial lawn. The number of contractions in the terminal bulb of pharynx was scored during a 30 seconds period immediately upon transfer using a stereomicroscope. This experiment was carried out in two independent biological replicates.

### Movement scoring/thrashing assay

Worms were grown as described and transferred to respective RNAi treatments at day 9. On day 20, 50 aged-synchronized worms from each knockdown condition were transferred individually in a 20 μL drop M9 buffer on a petri dish. After a 30 second recovery period, the numbers of body bends were scored during a 30 seconds period using a stereomicroscope. A body bend was defined as a change in the reciprocating motion of bending at the mid-body. This experiment was carried out in two independent biological replicates.

### Imaging and analysis of the axonal network

Worms were grown as described and transferred to respective RNAi treatments at day 9. On day 20, 100 aged-synchronized worms from each knockdown condition were paralyzed using 0.5% NaAz and mounted on a 2% agar pad on a glass microscope slide. GFP signal was acquired using a STED super-resolution microscope (Leica) at magnifications of 40x. Head regions of at least 100 different worms were imaged per knockdown condition and replicate. Axonal degeneration was scored on a three-point scale based on the amount of visible bubbling and neuron integrity. Scoring system: type 1 (intact axon without visible bubbling, kinks, or gaps); type 2 (slightly damaged axon, medium amount of bubbling, occasional kinks, very few to no gaps); type 3 (largely disrupted axon, high amount of bubbling, and frequent gaps). All images used for comparative quantifications were taken on the same day, with the same settings and by the same user. This experiment was carried out in two independent biological replicates.

### Western blotting

Worms were grown as described and transferred to respective RNAi treatments at day 9. For each time-point or knockdown condition: 200 synchronized animals were harvested, washed 3 times with M9 buffer and once with ddH_2_O, and suspended in 2x standard Laemmli buffer via 10 minute boiling. The samples were subjected to standard SDS-PAGE and western blotting. Antibodies used were monoclonal mouse anti-GFP (Roche,1:10,000), mouse monoclonal [6G6] anti-RFP (Chromotek,1:1000), mouse monoclonal anti-α-Tubulin (Sigma, 1:10,000), rabbit monoclonal anti-ubiquitin (lys48-specific Apu2 clone; Millipore, 1:1000). Antibodies were diluted in PBST (5% milk powder, 0.1% Tween20) which was also used as blocking agent. For the timecourse analysis of cleaved GFP and RFP, worms were maintained in liquid culture as described and a portion of the culture was harvested at each timepoint following a 40% Percoll wash. This wash was repeated in the event that the cleaned worms contained less than 95% living worms. Each western blot is representative for similar results obtained in two independent biological replicates.

### Chloroquine treatment

To pharmacologically inhibit the lysosome, animals were treated with chloroquine diphosphate salt (Sigma-Aldrich). Briefly, worms were grown as described in liquid culture until day 14 and subsequently incubated with 20 mM chloroquine or DMSO vehicle control for 24 hours. The drug treatment was performed in M9 liquid medium supplemented with EV HT115 bacteria at a density of 1x10^9^ cell ml^-1^ while shaking at 20 °C. The worms were harvested at day 15, following a 40% Percoll wash. This wash was repeated in the event that the cleaned worms contained less than 95% living worms.

### Statistics

All statistical analysis was performed with Graph Prism 6. p-values for lifespan curves were calculated using the Log-Rank (Mantel Cox) test. MTL was calculated as the mean remaining lifespan of the worms from the day of first treatment. p-values for quantification of GFP::LGG-1 foci were calculated using the non-parametric Mann Whitney U test. p-values for: muscle health, pharynx integrity, and neuronal integrity were calculated using a Chi-square test with two tails and 95% confidence interval. p-values for: qPCRs, body bend counts, pharynx pumping, and chymotrypsin assay were calculated using a two tailed t-test. Significance was scored as follows for all experiments: **** p<0.0001, *** p<0.001, ** p<0.01, * p<0.05.

## Acknowledgements

We thank IMB Core Facilities for their support, especially the Media Lab, the Microscopy and Bioinformatic Core Facility. Particular thanks to Kolja Becker for designing the Worm Count program for lifespan assays. We wish to thank Alicia Meléndez (Queens College, CUNY, USA) for providing us with the BEC-1::RFP reporter strain, and Scott Kennedy (Harvard Medical School, USA) for providing bacteria expressing dsRNA constructs targeting *GFP*. Some strains were provided by the CGC, which is funded by NIH Office of Research Infrastructure Programs (P40 OD010440). This work was supported by the Boehringer Ingelheim Foundation and a Marie Curie Reintegration Grant (321683, Call FP7-PEOPLE-2012-CIG). The Large Particle Sorter (Biosorter^®^, Union Biometrica) used for our large-scale RNAi longevity screen was financed through the DFG Major Research Instrumentation Program (INST 247/768-1 FUGG). We thank www.biographix.cz for the graphical abstract in Figure 4F.

